# Rewiring of transcription factor binding in differentiating human embryonic stem cells is constrained by DNA sequence repeat symmetry

**DOI:** 10.1101/375956

**Authors:** Matan Goldshtein, David B. Lukatsky

## Abstract

We analyze design principles of transcription factor (TF) recognition by genomic DNA in differentiating human embryonic stem cells for 36 TFs and five histone modifications in four developmental layers, using the data recently measured by Tsankov et al. This analysis reveals that DNA sequence repeat symmetry plays a central role in defining TF-DNA binding preferences across different developmental layers. In particular, we find that different TFs bind similar symmetry patterns within a given developmental layer. While the TF cluster content undergoes modifications upon transitions between different developmental layers, most TFs possess dominant preferences for similar DNA repeat symmetry types. Histone modifications also exhibit strong preferences for similar DNA repeat symmetry patterns, with the symmetry strength differentiating between different histone modifications. Overall, our findings show that despite the enormous sequence complexity of the TF-DNA binding landscape in differentiating human embryonic stem cells, this landscape can be quantitatively characterized in simple terms, using the notion of DNA sequence repeat symmetry.

## INTRODUCTION

The recently identified genome-wide TF-DNA binding landscape for 36 transcription factors (TFs) in differentiating human embryonic stem (ES) cells in four developmental layers demonstrates enormous complexity and plasticity of genomic DNA sequence recognition by TFs (1). In particular, these measurements identified TF-DNA binding preferences and the epigenetic landscape in the human ES cell line, HUES64 (ESc), and in four additional developmental layers Mesendoderm (dMS), Endoderm (dEN), Mesoderm (dME), and Ectoderm (dEC) (1).

The known specific TF binding motifs (TFBMs) populate a highly limited fraction of experimentally bound genomic sequences, and for many proteins, such as POL2, THAP11 and TRIM28 analyzed in this work, specific motifs have not been identified (1). For example, for two key transcription regulators, EOMES and OTX2, only 6-10% and 14-22% experimentally bound DNA sequences, respectively, contain specific TFBMs for these factors (1).

In recent years, it has been experimentally shown for many TFs, both *in vivo* and *in vitro*, that in addition to specific TFBMs, other factors, such as DNA shape (2-4) or certain non-consensus repetitive DNA sequence elements (5) can significantly affect TF-DNA binding. In general, it is now widely recognized that genomic DNA context outside of specific TFBMs significantly affects, and in many cases dominates, TF-DNA binding preferences (6-9).

In this work, we investigate DNA sequence repeat symmetries responsible for TF recognition in different developmental layers. Our central working hypothesis suggests that such repeat symmetries dominate TF-DNA binding preferences. Remarkably, we find that the majority of different TFs preferentially bind DNA sequences characterized by similar repeat symmetries. In order to quantitatively characterize DNA sequence repeat symmetries, we use one of the simplest possible measures for symmetry, namely, the nucleotide pair correlation function, developed and tested by us recently (10).

We have shown in the past using both biophysical modeling (5,10,11) and high-throughput *in vitro* measurements of TF-DNA binding preferences (5) that certain non-consensus, repetitive DNA sequence elements exert an entropy-dominated, statistical interaction potential on TFs. We use the term non-consensus TF-DNA binding free energy in order to describe this statistical interaction potential (5). Depending on the DNA symmetry type, the presence of certain repetitive DNA sequence elements can enhance or reduce the TF-DNA binding free energy (5). In the present work, we perform a genome-wide repetitive symmetry analysis that does not utilize any biophysical model, and does not have any fitting parameters. Experimentally identified genomic DNA sequences (TF binding peaks) constitute the only input to our simple computational procedure described below.

Here we identify the dominant DNA symmetry types that appear to influence large clusters of TFs in different developmental layers. In particular, we show that complex rewiring of TF-DNA binding network upon developmental transitions between different developmental layers can be quantified by the variation of the **DNA repeat symmetry type** and **DNA repeat symmetry strength** defined below.

## MATERIALS AND METHODS

### Computational procedure: Mapping DNA repeat symmetry

We analyze the micrococcal nuclease (MNase)-based ChIP-seq (MNChIP-seq) data for 36 TFs and five histone modifications in four developmental layers of differentiating human embryonic stem cells (1). All the data is publicly available at the Gene Expression Omnibus (GEO) under the accession number, GSE61475 (1). We analyze the DNA sequence repeat symmetry properties for each sequence set of TF binding peaks experimentally identified by MNChIP-seq (1).

We now define the measure for DNA sequence symmetry used to characterize genomic repetitive DNA sequence elements. Specifically, here we use the nucleotide pair-correlation function η_αα_(*x*), similar to the one used in our previous work (10). This correlation function, η_αα_(*x*), is proportional to the probability to find two nucleotides of the type *α* separated by the relative distance *x* along the genome, η_αα_(*x*)=(*N*_αα_(*x*)-<*N*_αα_(*x*)>_rand_)/*L*. For a given set of DNA sequences, *N*_αα_(*x*) is the total number of nucleotide pairs of the type *α* separated by the relative distance *x*, and <*N*_αα_(*x*)>_rand_ is the corresponding average number of nucleotide pairs in the randomized sequence set, and *L* is the total length of DNA sequences in the set. The randomization procedure randomly reshuffles each DNA sequence in the set, keeping the GC-content of each sequence intact. The averaging, <*N*_αα_(*x*)>_rand_, is performed with respect to ten random realizations of the original sequence set. Such randomization procedure normalizes the varying genomic GC-content, allowing us to compare symmetry properties of DNA sequences from different genomic locations characterized by a variable average GC-content.

For example, 60994 binding peaks were experimentally identified in human ES cells (ESc) for SRF protein (1). For each identified peak, we select 100-bp region in the middle of the peak. The entire set of these 60994 sequences is used in order to generate the correlation functions, η_αα_(*x*), for this protein in this developmental layer (Figure 1A). We use this procedure to generate the correlation functions for each TF or histone modification in each developmental layer. As a result, we obtain the entire set of η_αα_(*x*), for 36 TFs and five histone modifications in ES cells and in four developmental layers for all reported MNChIP-seq datasets (1). Although not all TFs and not all histone modifications were measured in each of four developmental layers, yet the existing data allows us to obtain a comprehensive, systems-level view on TF-DNA interaction network and on rewiring of this network in the course of ESc differentiation.

**Figure 1.**
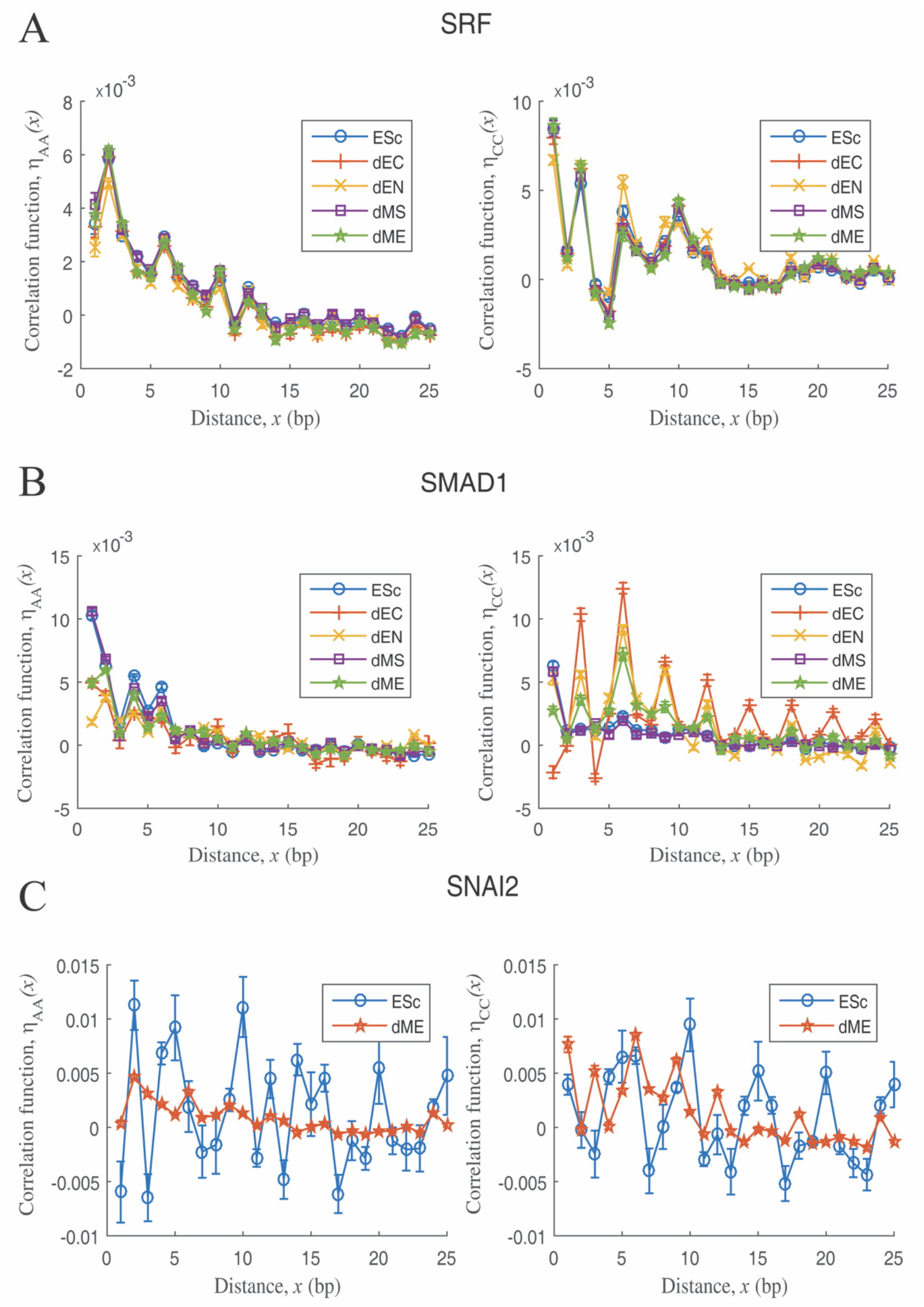
Correlation functions for the nucleotide spatial distribution identify DNA repeat symmetries selected by TFs in different developmental layers. The computed correlation function, *η_αα_*(*x*), for (A) SRF shows nearly identical symmetry type and symmetry strength in different developmental layers; (B) SMAD1 shows nearly identical symmetry type but varying symmetry strength in different developmental layers; (C) SNAI2 shows varying symmetry type and varying symmetry strength in different developmental layers. The correlation function, *η*_TT_(*x*), behaves similar to the shown *η*_AA_(*x*), and *η*_GG_(*x*) behaves similar to the shown *η*_CC_(*x*), Figure S1. In order to compute error bars, we divided each set of bound DNA sequences into five randomly chosen subgroups with equal number of sequences, and calculated *η_αα_*(*x*), for each subgroup. The error bars are defined as one standard deviation of *η_αα_*(*x*), between the subgroups.

## RESULTS

### DNA repeat symmetry type and DNA repeat symmetry strength

The computed correlation functions, η_AA_(*x*), η_TT_(*x*), η_CC_(*x*), and η_GG_(*x*), allow us to characterize the DNA sequence recognition specificity of TFs in terms of the **DNA repeat symmetry type** and the **DNA repeat symmetry strength** defined below. We stress the fact that the coordinate *x*, always represents the relative distance between the two nucleotides, and not the absolute distance with respect to a certain specific genomic location. For example, in the case of SRF (Figure 1A), η_CC_(*x*) and η_GG_(*x*), are characterized by the peaks at *x*=1, *x*=3, *x*=6, and *x*=10. The presence of such peaks means that DNA sequences bound by SRF in all developmental layers are statistically enriched in repetitive sequence patterns of the type [CC] (corresponding to *x*= 1), [CNNC] (corresponding to *x*=3), [CNNNNNC] (corresponding to *x*=6), *etc*., where N stands for any nucleotide type. On the contrary, η_AA_(*x*) and η_TT_(*x*), are characterized by the peaks at *x*=2, *x*=6, *etc*. It means that DNA sequences bound by SRF are also statistically enriched in repetitive sequence patterns of the type [ANA] (corresponding to *x*=2), [ANNNNNA] (corresponding to *x*=6), *etc*. (Figure 1A). Therefore, the relative positions of *x* characterized by the peaks in η_αα_(*x*), define the **DNA repeat symmetry type**. The height of the peaks (i.e. the magnitude of η_αα_(*x*) at a given peak position, *x*) define the **DNA repeat symmetry strength**.

### DNA repeat symmetry selection by TFs and histone modifications in developing human embryonic stem cells

We now aim to reveal a genome-wide view of DNA repeat symmetry selection by 36 TFs and five histone modifications in developing human ESc. Figure 1 shows examples of computed correlation functions, η_αα_(*x*), for three TFs in different developmental layers and in undifferentiated ESc. Here, we selected examples from three representative groups of TF (Figure 1). The first group corresponds to TFs (SRF) that show highly similar η_αα_(*x*), in all developmental layers (Figure 1A). The second group shows examples of TFs (SMAD1) with the similar DNA repeat symmetry type (i.e. similar peak positions), yet with the varying DNA repeat symmetry strength (i.e. varying peak heights) across different developmental layers (Figure 1B). The third group shows examples of TFs (SNAI2) with both the varying DNA repeat symmetry type and strength (Figure 1C).

The entire set of correlation functions computed for all 36 TFs in different developmental layers is shown in Figure S1.

The five measured histone modifications H3K27ac, H3K4me1, H3K4me3, H3K9me1, and H3K27me3 fall into the second group (i.e. similar DNA repeat symmetry type and varying symmetry strength across different developmental layers) (Figure 2). In particular, for the nucleotide types A and T, for all histone modifications (but H3K4me3 in the dMS developmental layer) the peaks in η_AA_(*x*) and η_TT_(*x*) are observed at *x*=2, *x*=4, *x*=6, *x*=8, *x*=10, *x*=12, and additional small peaks are observed at larger *x* (Figure 2). The only difference in the H3K4me3 histone modification is that it does not show a peak at *x*=2 (Figure 2). For C and G nucleotides, the peaks in η_CC_(*x*) and η_GG_(*x*) are observed at *x*=1, *x*=6, *x*=9, *x*=12, *x*=15 and even at larger *x* (Figure 2). Therefore, the histone DNA sequence specificity is achieved by varying the DNA repeat symmetry strength (peak heights), while keeping the DNA symmetry type (peak positions) practically fixed.

**Figure 2.**
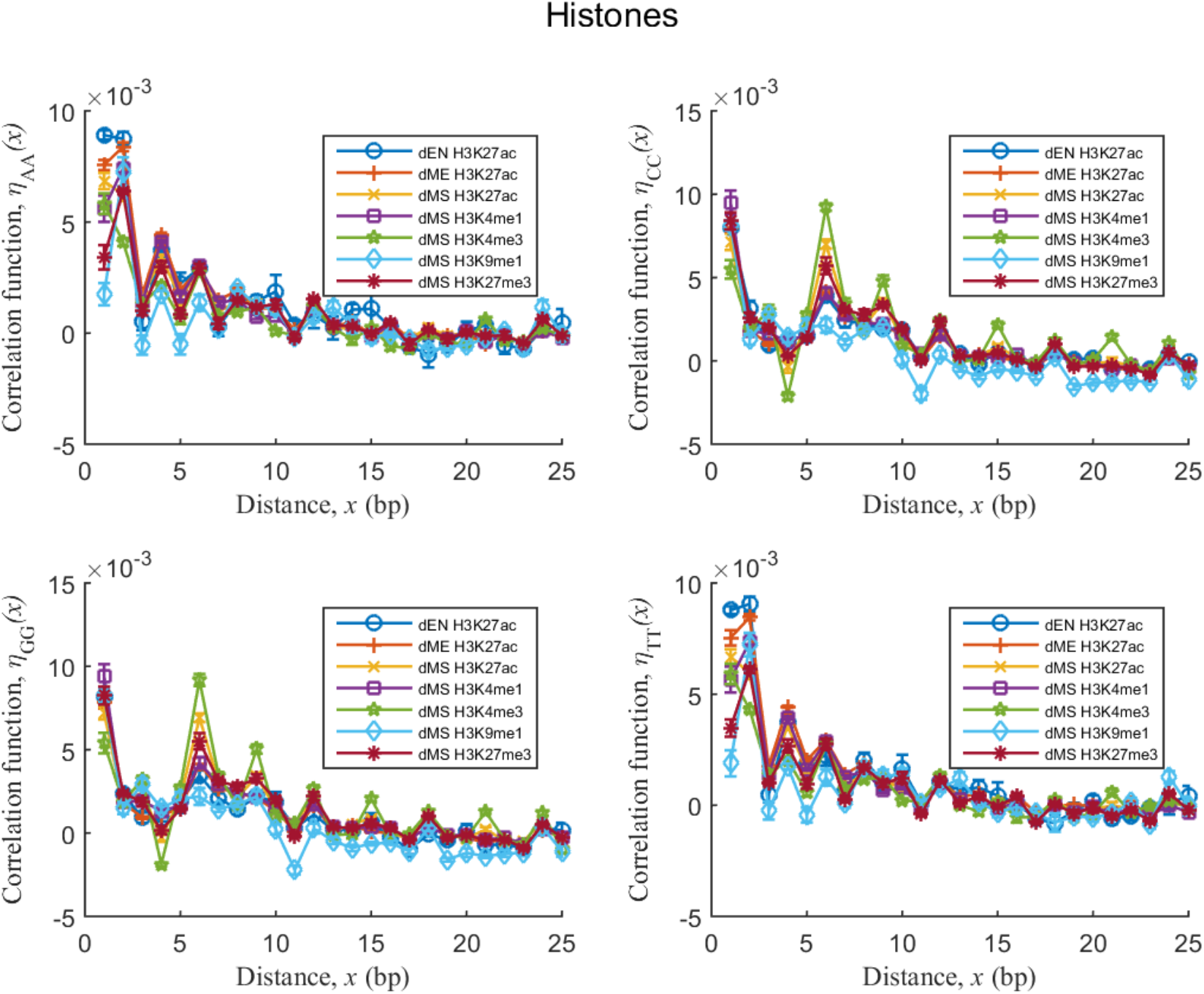
Correlation functions computed for different epigenetic histone modifications in different developmental layers show similar symmetry type (i.e. similar peak positions) but varying symmetry strength (i.e. varying peak heights). The error bars are defined similar to Figure 1.

In order to compare the degree of similarity between DNA repeat symmetry preferences of the entire set of TFs and histone modifications, we show the resulting heatmaps for all TF pairs for each developmental layer (Figure 3 and Figure S2). In order to construct the heatmap, for each pair of TFs in a given developmental layer, we compute the Pearson correlation coefficient, R, between the two correlation functions, η_αα_(*x*), characterizing the two TFs, separately for each nucleotide type α (Figure 3A and B). We perform this procedure for all pairs of TFs in a given developmental layer, and then represent the clustered distribution of the Pearson correlation coefficients, *R*, as a heatmap (Figure 3C-F). The most notable feature of the resulting heatmap is a significant degree of clustering between different TFs in the developmental layers (Figure 3 and Figure S2). For example, the six TFs (NANOG, OTX2, SMAD1, OCT4 (POU5F1), TCF4, and SOX2) reported as clustered in ESc and dMS (1), also appear almost entirely in one cluster in the heatmaps (all these TFs except OCT4 appear in one cluster in ESc, Figure 3D, and except SOX2 in dMS, Figure 3F).

**Figure 3.**
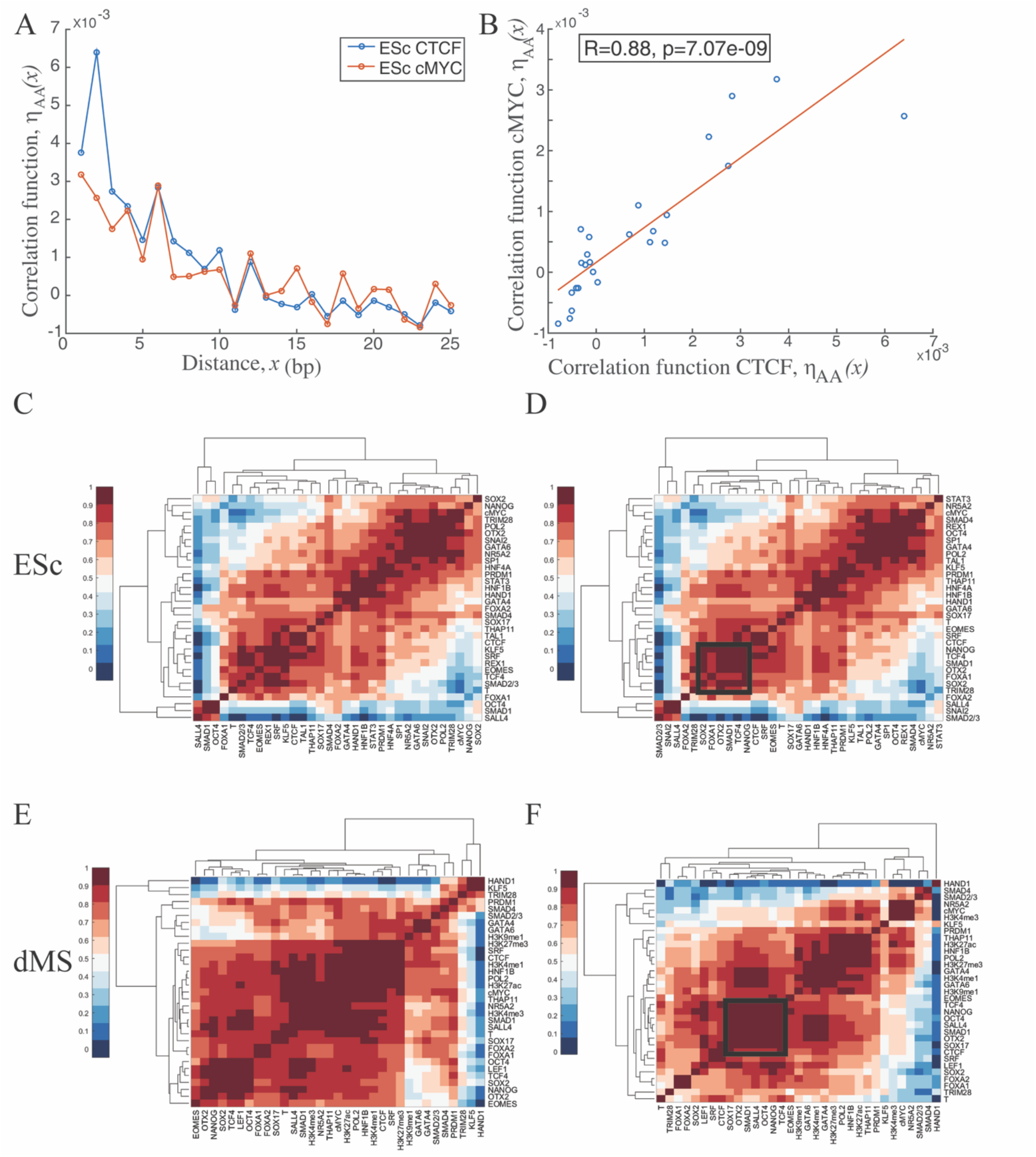
Correlation heatmaps characterizing the degree of similarity between the DNA repeat symmetry selected by TFs in a given developmental layer. (A) and (B) illustrate the definition of the computed Pearson correlation coefficient between each pair of TFs, using CTCF and cMYC as examples. (A) The computed correlation functions *η*_AA_(*x*), for CTCF and cMYC. (B) Computed linear correlation coefficient (the Pearson correlation coefficient) between *η*_AA_(*x*) for CTCF (x-axis) and cMYC (y-axis). Each point in this plot represents the value of *η*_AA_(*x*) for CTCF (x-coordinate) and cMYC (y-coordinate), respectively, for a given relative distance, *x*. The corresponding *p*-value is also shown. (C-F) Heatmaps generated by the Matlab *clustergram* function represent the computed Pearson correlation coefficients for all pairs of TFs and histone modifications for specific developmental layer, as follows: (C) ESc for nucleotide A, (D) ESc for nucleotide C, (E) dMS for nucleotide A, and (F) dMS for nucleotide C. Black frame emphasizes five out of six TFs (NANOG, OTX2, SMAD1, OCT4 (POU5F1), TCF4, and SOX2) reported as clustered in ESc and dMS (1), which also appear in one cluster (all these TFs except OCT4 appear in one cluster in ESc, and except SOX2 in dMS) in the heatmaps (D) and (F), respectively.

Even though the cluster content undergoes transformations upon developmental transitions between different developmental layers, a high degree of clustering is apparent in all developmental layers (Figure 3 and Figure S2). Such high degree of clustering stems from the fact that different TFs tend to preferentially bind certain DNA repeat symmetry types (i.e. DNA sequences characterized by similar peak positions in the correlation functions, η_αα_(*x*)) (Figure 4 and Figure S3). Therefore, although DNA binding preferences of each TF are characterized by a unique signature represented by the entire profile of η_αα_(*x*), statistically, on average, there is a high degree of similarity between many TFs with respect to the selected symmetry types.

**Figure 4.**
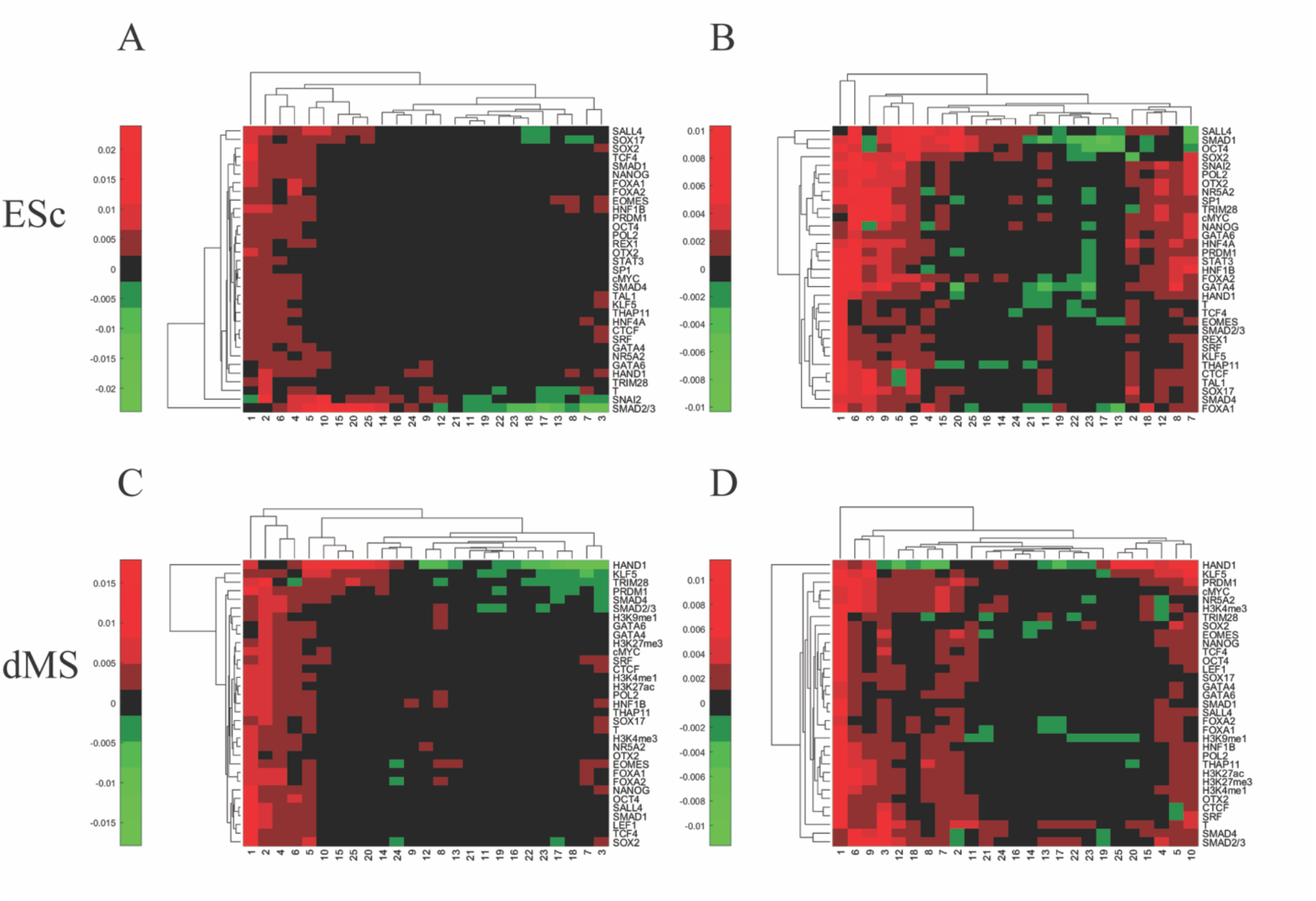
Clustered, heatmap representations of the computed correlation functions, *η*_αα_(*x*), for different TFs in a given developmental layer. TFs (y-axis) are clustered by similarities of the symmetry strength (i.e. the magnitude of *η*_αα_(*x*), represented by the shown color-code) at different relative distances, *x* (x-axis) in a given developmental layer. The heatmaps reveal, which symmetry types are more abundant in a given developmental layer for a given nucleotide type, as follows: (A) ESc for nucleotide A, (B) ESc for nucleotide C, (C) dMS for nucleotide A, and (D) dMS for nucleotide C.

### Protein-protein interactions dramatically modify DNA repeat symmetry selection by SMAD1

Using SMAD1 protein as a case study, we now ask the question how protein-protein interactions modify DNA repeat symmetry selection by TFs in different developmental layers. In a series of MNChIP-seq experiments performed by Tsankov et al. (1), genome-wide binding preferences of SMAD1 were measured in different developmental layers under suppressed expression of its key binding partner, GATA4 (1). Strikingly, we observe a dramatic change in the DNA repeat symmetry selection by SMAD1 upon silencing of GATA4 (Figure 5). In particular, upon silencing of GATA4, η_CC_(*x*) and η_GG_(*x*) in the dEN and dME germ layers (later developmental stages), behave similar to η_CC_(*x*) and η_GG_(*x*) in the dMS and ESc developmental layers (earlier developmental stages), at normal expression levels of GATA4 (Figure 5 and Figure 1B). Interestingly, despite the fact that SMAD1 recruits histone acetyltransferase, binding preferences of H3K27ac are not noticeably affected by the silencing of GATA4 (Figure S4). We conclude therefore, that protein-protein interactions between TFs can significantly modify the DNA repeat symmetry selection by a given TF in different developmental layers. However, at least based on the current example of SMAD1, the modified DNA symmetry selected by SMAD1 (at suppressed GATA4) is highly restricted by the DNA symmetry it selects in other developmental layers in normal conditions (i.e. at wild-type expression levels of GATA4). Such symmetry selection switching, modulated by protein-protein interactions might represent a general mechanism regulating TF-DNA binding preferences in ESc, and it would be thus important to experimentally test additional pairs of interacting TFs. In their recent, seminal study Fiore and Cohen developed a high-throughput method for the *in vivo* testing of the effect of the interaction between TFs (12). Using mouse embryonic stem cells as a model system, Fiore and Cohen demonstrated that the interaction between the key pluripotency modulators, such as POU5F1 (OCT4), SOX2, KLF4, and ESRRB, indeed significantly affects the gene expression (12). Their method involving the construction of synthetic promoter library of *cis*-regulatory elements (CREs), can be adopted to test the effect of protein-protein interactions in the presence of repetitive DNA sequence elements predicted to affect certain TFs.

**Figure 5.**
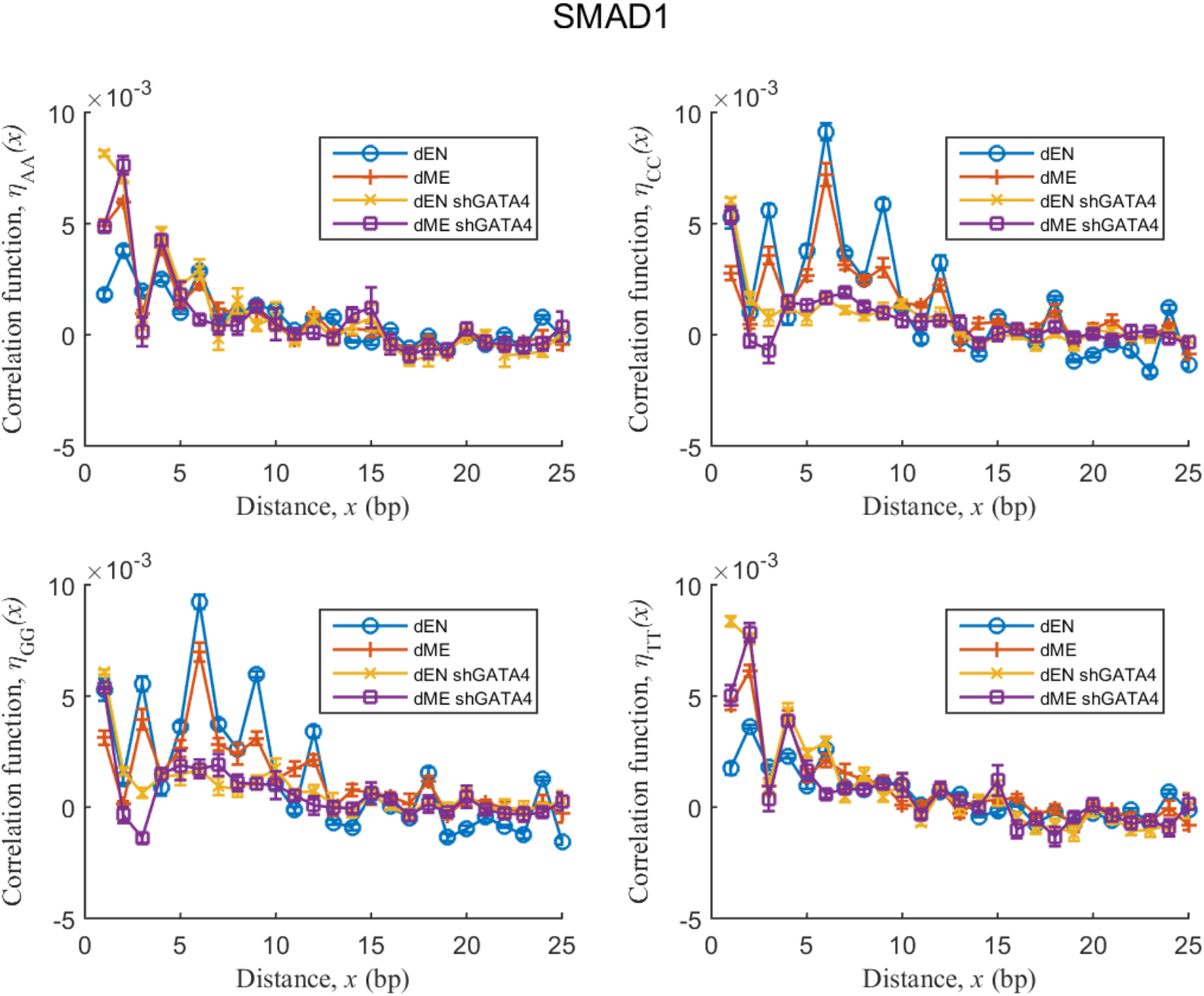
The correlation functions, *η_αα_*(*x*), computed for DNA sequences bound by SMAD1, measured in cells with silencing of GATA4 (designated shGATA4) compared to wild type cells (i.e. at normal expression levels of GATA4). The error bars are defined similar to Figure 1.

## DISCUSSION

Here we show that the DNA sequence repeat symmetry represents an important sequence signature providing the specificity for TF binding in developing human ESc. In particular, binding preferences of TFs towards genomic DNA can be quantitatively characterized by the DNA repeat symmetry type and the DNA repeat symmetry strength. The former parameter (symmetry type) corresponds to the positions of the peaks of the correlation functions η_AA_(*x*), η_TT_(*x*), η_CC_(*x*), and η_GG_(*x*), with respect to the relative distance, *x*, between the two given nucleotides. The latter parameter (symmetry strength) corresponds to the magnitude of the heights of these correlation functions at particular peak positions, *x*. The correlation functions, η_αα_(*x*), are defined in such a way that different genomic positions can be quantitatively compared despite the fact that the average GC-content is varying along the genome.

The most striking conclusion that emerges from the DNA repeat symmetry analysis of bound genomic regions is the fact that despite the enormous sequence complexity of the TF-DNA binding landscape, only few DNA repeat symmetry types are selected by the majority of TFs in all developmental layers (Figure 4 and Figure S3). For example, the correlation functions η_AA_(*x*) and η_TT_(*x*), for the majority of TFs are peaked at *x*=1, *x*=2, *x*=4, and *x*=6 (Figure 4 and Figure S3). The correlation functions, η_CC_(*x*) and η_GG_(*x*), for the majority of TFs show peaks at *x*=1, *x*=3, *x*=6, *x*=9, and *x*=12 (Figure 4 and Figure S3). In addition, yet to a lesser degree, additional repeating patterns are observed at *x*=5, *x*=10, for the C and G nucleotide types (Figure 4 and Figure S3). This means that statistically, on average, only few dominant repetitive DNA sequence patterns significantly contribute to establishing the genome-wide TF-DNA binding profile. However, despite the observed similarity, DNA binding preferences of each TF are characterized by a unique signature represented by the entire profile of η_αα_(*x*), i.e., the entire profile of the DNA symmetry type (peak positions along *x*) and the DNA symmetry strength (peak heights).

The DNA repeat sequence symmetry selection by five different histone modifications in different developmental layers, represents an example of striking universality (robustness) in the DNA repeat symmetry type (i.e. peak positions) (Figure 2 and Figure S4). The specificity for different histone modifications towards genomic DNA in human embryonic stem cells is thus tuned by tuning the DNA repeat symmetry strength (i.e. peak heights), while keeping the DNA repeat symmetry type (i.e. peak positions) nearly invariant for all histone modifications (Figure 2).

Overall, our findings thus suggest that DNA repeat symmetry significantly contributes towards establishing the genome-wide TF-DNA binding profile in developing human embryonic stem cells.

The central question that needs to be discussed now is what molecular mechanism responsible for the observed genome-wide dependence of TF binding preferences on DNA repeat symmetry? We have suggested in the past, and validated experimentally for a number of TFs, that certain repetitive DNA sequence elements exert the entropy-dominated TF-DNA binding free energy landscape (5,10,11,13). The main difficulty here stems from the fact that in human ESc (as well as in any other type of cells) there are additional factors, besides the DNA sequence alone, that influence TF-DNA binding preferences. Such factors include, first of all, protein-protein interactions affected by protein expression levels, DNA and histone epigenetic modifications, and chromatin folding affected by itself by a number of epigenetic factors (12,14-20). Our analysis of the effect of protein-protein interactions on the DNA repeat symmetry selection by TFs, using SMAD1 interacting with GATA4 as a case study, shows that such interactions can dramatically affect DNA repeat symmetries selected by TFs (Figure 5). Direct experimental validation of the observed DNA repeat symmetry effect on genome-wide TF-DNA binding preferences in developing human ESc, should constitute a future step towards better mechanistic understanding of the findings presented in this work. Such validation can be performed by cloning DNA sequences enriched with particular repeat symmetries (predicted to bind certain TFs) into a selected set of promoters and enhancers and measuring the resulting binding preferences of these TFs. A high-throughput method, recently developed by Fiore and Cohen, using a synthetic promoter library of cis-regulatory elements, might constitute a suitable model system for this purpose (12).

## ACKNOWLEDGEMENTS

We acknowledge the generous support and professional help of Prof. Smadar Cohen and the Regenerative Medicine and Stem Cell (RMSC) Research Center at Ben-Gurion University of the Negev.

## FUNDING

None declared.

## CONFLICT OF INTEREST

None declared.

## REFERENCES

1. Tsankov, A.M., Gu, H., Akopian, V., Ziller, M.J., Donaghey, J., Amit, I., Gnirke, A. and Meissner, A. (2015) Transcription factor binding dynamics during human ES cell differentiation. Nature, 518, 344–349.

2. Gordan, R., Shen, N., Dror, I., Zhou, T., Horton, J., Rohs, R. and Bulyk, M.L. (2013) Genomic regions flanking E-box binding sites influence DNA binding specificity of bHLH transcription factors through DNA shape. Cell reports, 3, 1093–1104.

3. Slattery, M., Zhou, T., Yang, L., Dantas Machado, A.C., Gordan, R. and Rohs, R. (2014) Absence of a simple code: how transcription factors read the genome. Trends in biochemical sciences, 39, 381–399.

4. Rossi, M.J., Lai, W.K.M. and Pugh, B.F. (2018) Genome-wide determinants of sequence-specific DNA binding of general regulatory factors. Genome research, 28, 497–508.

5. Afek, A., Schipper, J.L., Horton, J., Gordan, R. and Lukatsky, D.B. (2014) Protein-DNA binding in the absence of specific base-pair recognition. Proceedings of the National Academy of Sciences of the United States of America, 111, 17140–17145.

6. Xin, B. and Rohs, R. (2018) Relationship between histone modifications and transcription factor binding is protein family specific. Genome research.

7. Le, D.D., Shimko, T.C., Aditham, A.K., Keys, A.M., Longwell, S.A., Orenstein, Y. and Fordyce, P.M. (2018) Comprehensive, high-resolution binding energy landscapes reveal context dependencies of transcription factor binding. Proceedings of the National Academy of Sciences of the United States of America, 115, E3702–e3711.

8. Esadze, A., Kemme, C.A., Kolomeisky, A.B. and Iwahara, J. (2014) Positive and negative impacts of nonspecific sites during target location by a sequence-specific DNA-binding protein: origin of the optimal search at physiological ionic strength. Nucleic acids research, 42, 7039–7046.

9. Pugh, B.F. and Venters, B.J. (2016) Genomic Organization of Human Transcription Initiation Complexes. PloS one, 11, e0149339.

10. Goldshtein, M. and Lukatsky, D.B. (2017) Specificity-Determining DNA Triplet Code for Positioning of Human Preinitiation Complex. Biophysical journal, 112, 2047–2050.

11. Sela, I. and Lukatsky, D.B. (2011) DNA sequence correlations shape nonspecific transcription factor-DNA binding affinity. Biophysical journal, 101, 160–166.

12. Fiore, C. and Cohen, B. (2016) Interactions between pluripotency factors specify cis-regulation in embryonic stem cells. Genome research, gr. 200733.200115.

13. Imashimizu, M., Afek, A., Takahashi, H., Lubkowska, L. and Lukatsky, D.B. (2016) Control of transcriptional pausing by biased thermal fluctuations on repetitive genomic sequences. Proceedings of the National Academy of Sciences of the United States of America, 113, E7409–e7417.

14. Eberharter, A. and Becker, P.B. (2002) Histone acetylation: a switch between repressive and permissive chromatin: Second in review series on chromatin dynamics. EMBO Reports, 3, 224–229.

15. Javaid, N. and Choi, S. (2017) Acetylation-and Methylation-Related Epigenetic Proteins in the Context of Their Targets. Genes, 8, 196.

16. Quina, A.S., Buschbeck, M. and Di Croce, L. (2006) Chromatin structure and epigenetics. Biochemical pharmacology, 72, 1563–1569.

17. Verdone, L., Agricola, E., Caserta, M. and Di Mauro, E. (2006) Histone acetylation in gene regulation. Briefings in Functional Genomics, 5, 209–221.

18. Rotem, A., Ram, O., Shoresh, N., Sperling, R.A., Goren, A., Weitz, D.A. and Bernstein, B.E. (2015) Single-cell ChIP-seq reveals cell subpopulations defined by chromatin state. Nature biotechnology, 33, 1165–1172.

19. Teif, V.B. and Cherstvy, A.G. (2016) Chromatin and epigenetics: current biophysical views. AIMS Biophysics, 3, 88–98.

20. de Dieuleveult, M., Yen, K., Hmitou, I., Depaux, A., Boussouar, F., Bou Dargham, D., Jounier, S., Humbertclaude, H., Ribierre, F., Baulard, C. et al. (2016) Genome-wide nucleosome specificity and function of chromatin remodellers in ES cells. Nature, 530, 113–116.

